# Interleukin-17 regulates neuron-glial communications, inhibitory synaptic transmission and neuropathic pain after chemotherapy

**DOI:** 10.1101/344945

**Authors:** Hao Luo, Hui-Zhu Liu, Xin Luo, Sangsu Bang, Zi-Long Wang, Gang Chen, Ru-Rong Ji, Yu-Qiu Zhang

## Abstract

The proinflammatory cytokine Interleukin-17 (IL-17) is produced mainly by Th17 cells and has been implicated in pain regulation. However, synaptic mechanisms by which IL-17 regulates pain transmission are unknown. Here we report that glia-produced IL-17 suppresses inhibitory synaptic transmission in spinal cord pain circuit and drives chemotherapy-induced neuropathic pain. We observed respective expression of IL-17 and its receptor IL-17R in spinal cord astrocytes and neurons. Patch clamp recording in spinal cord slices revealed that IL-17 not only enhanced EPSCs but also suppressed IPSCs and GABA-induced currents in lamina II_o_ somatostatin-expressing neurons. Spinal IL-17 was upregulated after paclitaxel treatment, and intrathecal IL-17R blockade reduced paclitaxel-induced neuropathic pain. In dorsal root ganglia, respective IL-17 and IL-17R expression in satellite glial cells and neurons was sufficient and required for inducing neuronal hyperexcitability after paclitaxel. Together, our data show that IL-17/IL-17R mediate both central and peripheral neuron-glial interactions in chemotherapy-induced peripheral neuropathy.

## Introduction

Pro-inflammatory cytokines such as TNF-α, IL-1β, and IL-18 play important roles in the pathogenesis of chronic pain (Sommer, 1999, Zelenka et al., 2005, Milligan et al., 2001, Yang et al., 2015, Miyoshi et al., 2008, Sweitzer et al., 1999, Guo et al., 2007). Increasing evidence suggests that glial cells such as microglia and astrocytes are activated in pathological pain conditions to produce these pro-inflammatory cytokines. Especially, these cytokines act as neuromodulators and regulate pain via neuron-glial interactions (Ji et al., 2013, Grace et al., 2014). Compared to TNF and IL-1β, and IL-6, much less is known about the role of IL-17 in pain regulation. IL-17, referred to as IL-17A in the literature, is a proinflammatory cytokine produced by Th17 cells (Miossec and Kolls, 2012, Korn et al., 2009). The IL-17 family consists of six ligands (IL-17A–F) and five receptors (IL-17RA–IL-17RE) in mammals, (Gaffen, 2009). IL-17 was shown to regulate rheumatoid arthritis and immune responses by increasing the production of IL-6 and IL-8(Hwang et al., 2004). Binding of IL-17 to its receptor (IL-17RA) induces the activation of nuclear factor-κB (NF-κB) via ACT1 and TNF receptor-associated factor 6 (TRAF6) in rheumatoid arthritis(Hot and Miossec, 2011). However, little is known about non-transcriptional regulation of IL-17.

Recently, IL-17 was found to regulate inflammatory responses associated with neuropathic pain induced by nerve injury. IL-17 levels are upregulated in the injured nerves in neuropathic pain models(Noma et al., 2011, Kleinschnitz et al., 2006). IL-17 receptor (IL-17R) was detected in most neurons in dorsal root ganglion (DRG) as well as in cultured DRG neurons (Segond von Banchet et al., 2013, Richter et al., 2012). IL-17A-deficient mice showed less mechanical hyperalgesia compared to normal mice after zymosan injection(Segond von Banchet et al., 2013) or partial ligation of the sciatic nerve (Kim and Moalem-Taylor, 2011). Further, Intraplantar (Kim and Moalem-Taylor, 2011, McNamee et al., 2011) or intra-knee (Pinto et al., 2010) injection of recombinant IL-17 is sufficient to induce hyperalgesia. Notably, IL-17 can also be produced by spinal cord astrocytes, and astrocytic IL-17 may play a role in inflammatory pain (Meng et al., 2013). A recent study found that physiological levels of IL-17 can act directly on interneurons to increase their responsiveness to presynaptic input (Chen et al., 2017). Despite these previous studies, it remains elusive how IL-17 modulates spinal synaptic transmission in the pain circuit.

Chemotherapy-induced peripheral neuropathy (CIPN) is a common dose-limiting adverse effect and results in high incidence of neuropathic pain (Sisignano et al., 2014). There is evidence that spinal astrocytes but not microglia play an important role in the pathogenesis of paclitaxel-induced neuropathy (Zhang et al., 2012a, Luo et al., 2017). CIPN enhances excitability of primary sensory neurons associated with altered gene expression of neuronal ion channels. (Zhang and Dougherty, 2014) and also increases excitatory synaptic transmission in spinal cord substantia gelatinosa neurons (Li et al., 2015a).

The somatostatin-positive (SOM^+^) neurons are a subset of interneurons in the dorsal horn. There neurons are predominantly excitatory and express the vesicular glutamate transporter VGLUT_2_, a marker for glutamatergic excitatory neurons (Duan et al., 2018, Xie et al., 2018). Recently, Duan et al. demonstrate that SOM^+^ neurons are required to sense mechanical pain (Duan et al., 2014b). These neurons form a pain circuit by receiving input from capsaicin-sensitive C-fibers and sending output to lamina I projection neurons (Todd, 2010, Braz et al., 2014). SOM^+^ also receive input from inhibitory neurons (Duan et al., 2014b). Furthermore, these neurons exhibit remarkable plastic changes after inflammation and nerve injury and respond to inflammatory mediators (Park et al., 2011, Xie et al., 2018, Xu et al., 2010). Here, we investigated how IL-17 and IL-17R modulate excitatory and inhibitory synaptic transmission of SOM^+^ excitatory neurons in the normal and pathological pain conditions and further tested the involvement of IL-17/IL-17R signaling in paclitaxel-induced neuropathic pain model. Our findings demonstrate that IL-17 signaling contributes to paclitaxel-induced mechanical allodynia and dysregulations of excitatory and inhibitory synaptic transmission in spinal SOM^+^ neurons. Moreover, we reveal new insights into neuron-glial interactions in the spinal cord and DRG, by which IL-17 produced by astroglia or satellite glia enhance neuronal activities and excitability to promote neuropathic pain.

## Methods

### Animals

Most experiments were performed on adult C57BL/6 mice (8-10 weeks, male, purchased from Charles River). Some electrophysiology experiments were conducted in transgenic C57BL/6 mice (5-6 weeks). These mice express tdTomato fluorescence in somatostatin (SOM^+^) neurons, after Som-Cre mice were crossed with tdTomato Cre-reporter mice (Rosa26-floxed stop tdTomato mice), both from Jackson Laboratory, to generate conditional transgenic mice that express tdTomato in SOM^+^ neurons. All the animal procedures were approved by the Institutional Animal Care & Use Committee (IACUC) of Duke University and Fudan University.

Intraperitoneal (i.p.) injection of paclitaxel (PAX, 6 mg/kg for a single injection or 2 mg/kg for multiple injections at days 0, 2, 4, and 6) was given to generate chemotherapy-associated neuropathic pain (20). 7 days following the injection, spinal dorsal horns and CSF were collected.

### Reagents and drug injection

We purchased the recombinant mouse IL-17A protein (R&D System Inc., MN, USA., 421-ML), mouse IL-17 receptor A (IL-17 RA or IL-17R) antibody (R&D System Inc., MN, USA., MAB4481) and control IgG (R&D Systems Inc., MN, USA.). GABA, and glycine were obtained from Sigma-Aldrich. IL-17 was prepared as 1000 fold stock solution in 4 mM HCl and finally used at the concentration of 10 ng/mL. All compounds were prepared in artificial cerebrospinal fluid (ASCF). Picrotoxin, strychnine, AP-5 or CNQX were purchased from Sigma Company.

IL-17 or vehicle was delivered to CSF space between L5 and L6 vertebrae via a spinal cord puncture, which is made by a 30 Gage needle. Before puncture, the mice’ heads were covered by a piece of cloth. Ten microliters of solution were injected with a microsyringe. A successful spinal puncture was confirmed by a brisk tail-flick.

### ELISA

ELISA was performed using CSF and spinal cord tissues. The tissues were homogenized in a lysis buffer containing protease and phosphatase inhibitors (Sigma Chemical Co), and tissue samples were centrifuged (12,500×g for 10 min) to obtain extract proteins. CSF was collected from the cisterna magna. For each ELISA assay, 50 μg proteins, or 5 μL of CSF were used. ELISA was conducted according to manufacturer’s instructions (R&D Systems Inc., MN, USA., Cat# PM1700) and the standard curve was included in each experiment.

### Behavior

Animals were habituated to the testing environment for at least 2 days before the testing. Animals were kept in boxes on an elevated metal mesh floor. Mechanical allodynia was assessed by measuring paw withdrawal thresholds in response to a series of von Frey hairs (0.16-2.0 g, Stoelting Company). The withdrawal threshold was determined using the Dixon’s up-down method.

### Immunohistochemistry

Mice were deeply anesthetized with urethane and were transcardially perfused with normal saline followed by 4% paraformaldehyde in 0.1 M PB. The L4–L6 segments of the spinal cord were removed and postfixed for 24 h at 4°C, and then dehydrated in gradient sucrose at 4°C. Transverse spinal cord sections (30 um) were cut on a cryostat (model 1900, Leica). The sections were blocked with PBS containing 10% donkey serum and 0.3% Triton X-100 for 2 h at RT and then incubated for 48 h at 4°C with a mixture of rabbit anti-IL-17 (1:50) and mouse anti-NeuN (1:2000)/rabbit anti-IBA-1 (1:500)/mouse anti-GFAP (1:2000) antibodies, or rabbit anti-IL-17R (1:200) and anti-NeuN/GFAP/IBA-1. The sections were then incubated with a mixture of FITC-conjugated secondary antibodies (1:200; Jackson ImmunoResearch) for 2h at RT. Negative control was included by the omission of the primary antibodies. The stained sections were observed and the images captured with a confocal laser-scanning microscope (model FV1000, Olympus).

### Preparation of spinal cord slices and whole-cell patch-clamp recordings

The L4–L5 lumbar spinal cord segment was rapidly removed under urethane anesthesia (1.5 - 2.0 g/kg, i.p.) and transferred to ice-cold cutting ACSF containing (in mM) NaCl 80, KCl 2.5, NaH_2_PO_4_ 1.25, CaCl_2_ 0.5, MgCl_2_ 3.5, NaHCO_3_ 25, sucrose 75, ascorbate1.3, sodium pyruvate 3.0, oxygenated with 95% O_2_ and 5% CO_2_, pH 7.4. Transverse slices (450 μm) were cut on a vibrating blade microtome (Leica VT1200 S) and incubated in recording ACSF oxygenated with 95% O_2_ and 5% CO_2_ for at least 1 h at 32°C before recording. Slices were then transferred to the chamber and perfused with recording solution at a rate of 3 ml/min at RT. The recording ACSF contains the following (in mM): NaCl 125, KCl 2.5,CaCl _2_ 2, MgCl _2_ 1, NaH_2_PO_4_ 1.25, NaHCO_3_ 26, D-glucose 25.

The whole-cell patch clamp recordings were performed in lamina IIo SOM^+^ neurons in voltage-clamp mode. Patch pipettes (5–10 MΩ) were made of borosilicate glass on a horizontal micropipette puller (P-97, Sutter Instruments) analysis. For spontaneous excitatory postsynaptic currents (sEPSCs) recordings, pipette solution contained (in mM): potassium gluconate 120, KCl 20, MgCl_2_ 2, Na_2_ATP 2, NaGTP 0.5, HEPES 20, EGTA 0.5, adjusted to pH 7.3 with KOH. For spontaneous inhibitory postsynaptic currents (sIPSCs), pipette solution contained (in mM): CsCl 130, NaCl 9, MgCl2 1, EGTA 10, HEPES 10, adjusted to pH 7.3 with CsOH. After establishing the whole-cell configuration, neurons were held at −70 mV to record sEPSCs in the presence of 100 μM picrotoxin and 2 μM strychnine. Signals were filtered at 2 kHz and digitized at 5 kHz. NMDA receptor mediated EPSC was evoked by electrical stimulation of Lissauer’s tract, using a low Mg^2+^ recording ACSF (2.5 mM Ca^2+^, 0.25 mM Mg^2+^) with CNQX (10 μM), BMI, (10 μM) and strychnine (2 μM). A constant current pulse (0.3–0.5mA) at 0.05 Hz was applied to the Lissauer’s tract to evoke EPSC. When recording NMDA-EPSCs, a holding potential at −40 mV was used as indicated. GABA current and glycine current were induced by 100 μM GABA and 1 mM glycine, respectively. Data were collected with pClamp 10.1 software and analyzed with Mini Analysis and Clampfit.

### Whole-cell patch clamp recordings in dissociated mouse DRG neurons

DRGs were aseptically removed from 5-8 week-old mice and digested with collagenase (0.2 mg/ml, Roche)/dispase-II (3 mg/ml, Roche) for 120 min. Cells were placed on glass cover slips coated with poly-D-lysine and grown in a neurobasal defined medium (10% fetal bovine serum and 2% B27 supplement) at 37°C with 5% CO_2_ for 24 h before experiments.

### Human DRG neuron cultures and whole-cell patch clamp recordings

Non-diseased human DRGs were obtained from donors through National Disease Research Interchange (NDRI) with permission from the Duke University Institutional Review Board (IRB). Postmortem L3–L5 DRGs were dissected from donors and delivered in ice-cold culture medium to the laboratory at Duke University within 24–72 h of the donor’s death. Upon the delivery, DRGs were rapidly dissected from nerve roots and minced in a calcium-free HBSS (Gibco). Human DRG cultures were prepared as previously reported (Han et al., 2016, Chang et al., 2018). DRGs were digested at 37 °C in a humidified O2 incubator for 120 min with collagenase Type II (Worthington, 290 units/mg, 12 mg/ml final concentration) and dispase II (Roche, 1 unit/mg, 20 mg/ml) in PBS with 10 mM HEPES, pH adjusted to 7.4 with NaOH. hDRGs were mechanically dissociated using fire-polished pipettes, filtered through a 100 μm nylon mesh and centrifuged for 5 min (500g). The DRG cell pellet was resuspended, plated on 0.5 mg/ml poly-D-lysine–coated glass coverslips. DRG cultures were grown in Neurobasal medium supplemented with 10% FBS, 2% B-27 supplement, and 1% penicillin/streptomycin.

Whole-cell patch-clamp recordings in small-diameter (<55 μm) human DRG neurons were conducted at room temperature. We used an Axopatch-200B amplifier with a Digidata 1440A (Axon Instruments) to measure action potentials and resting membrane potential. The patch pipettes were pulled from borosilicate capillaries (World Precision Instruments, Inc.). The resistance of the pipettes was 3-4 MΩ, when filled with the pipette solution. The recording chamber (300 µl) was continuously superfused at the flow rate of 1-2 ml/min. Series resistance was compensated (> 80%) and leak subtraction was performed. Data were low-pass-filtered at 2 KHz and sampled at 10 KHz. The pClamp10.6 (Axon Instruments) software were used during experiments and Clampfit 10.6 were used for analysis. The pipette solution contained (in mM): potassium gluconate 126, NaCl 10, MgCl_2_ 1, EGTA 10, Na-ATP 2 and Mg-GTP 0.1, adjusted to pH 7.3 with KOH. The external solution contained: NaCl 140, KCl 5, CaCl_2_ 2, MgCl_2_ 1, HEPES 10, glucose 10, adjusted to pH 7.4 with NaOH. In current-clamp experiments, the action potentials were evoked by a current injection. The resting membrane potential was measured without a current injection.

### Data analysis and statistics

All data were expressed as mean ± S.E.M. ELISA and behavioral data were analyzed using Student’s t-test (two groups) or two-way ANOVA followed by post-hoc Bonferroni test. Electrophysiological data were tested using two-way ANOVA followed by post-hoc Bonferroni test or two-tailed paired t-test. The criterion for statistical significance was p < 0.05.

## Results

### Distinct cellular localization of IL-17 and IL-17R in spinal dorsal horn

As the first step to define the role of IL-17 and IL-17R in regulating spinal cord synaptic transmission and CIPN, we examined cellular location of IL-17 and IL-17R in spinal cord. Double immunofluorescence labeling demonstrated that IL-17 immunoreactivity (IR) was primarily colocalized with the astrocyte marker GFAP but not with the neuronal marker NeuN or the microglia marker IBA1 in spinal dorsal horn (Fig. 1A-C). Interestingly, IL-17R-IR showed distinct expression pattern. IL-17 receptor was predominantly colocalized with NeuN but not with GFAP or IBA1 in the spinal cord dorsal horn (Fig. 2A-C). The data indicate that IL-17 is expressed by astrocytes but its receptor is expressed by neurons in spinal cord. Notably, the expression of IL-17 and IL-17R was enriched in the superficial dorsal horn, where nociceptive input (C- and Aδ-afferents (Basbaum et al., 2009). These unique expression patterns of the ligand-receptor pair provide an anatomical base for IL-17 to mediate neuron-glial interaction in the spinal cord pain pathway.

**Figure 1.**
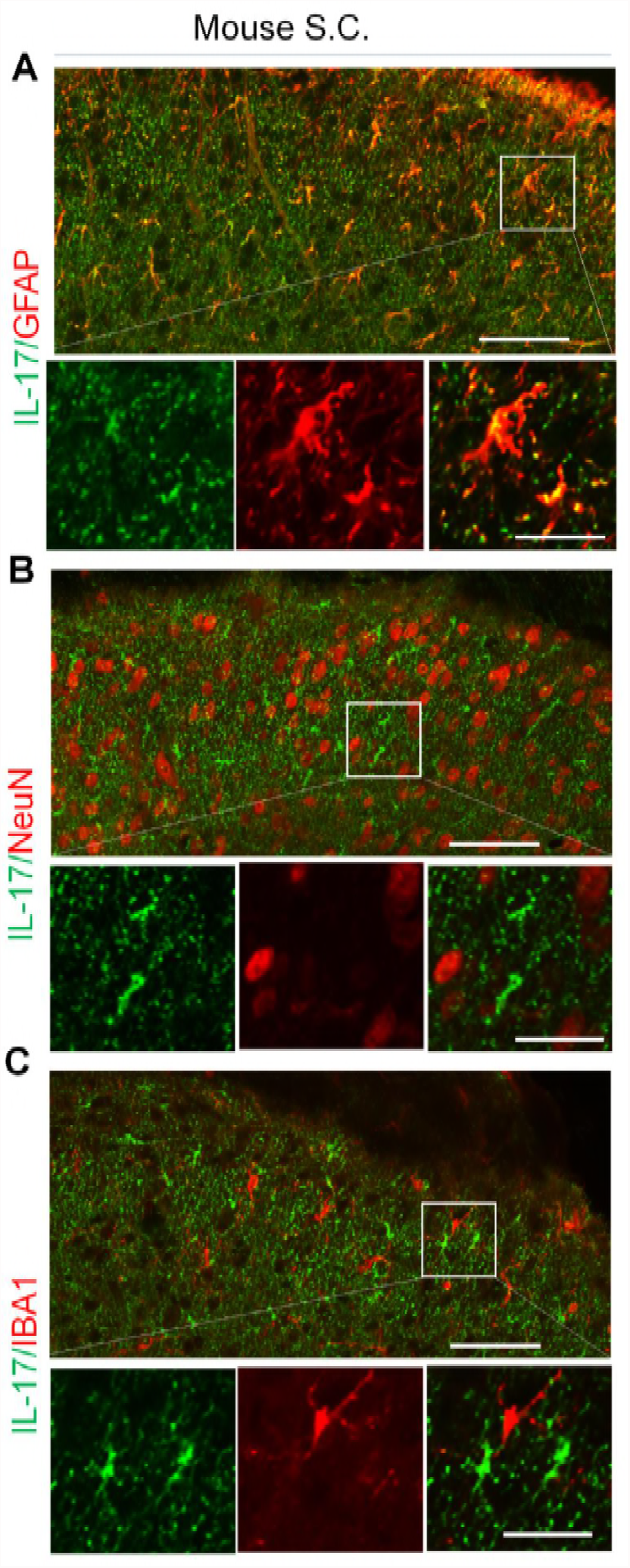
Photomicrographs showing IL-17 expression and colocalization of IL-17 with GFAP in spinal dorsal horn (SDH) Double labeling of IL-17 with astrocyte marker GFAP (**A**), neuron marker NeuN (**B**), and microglia marker IBA1 (**C**) in SDH. The white square in the top image is enlarged in three separate boxes with single and merged images in each picture (**A**, **B** and **C**). The scale bars represent 50 µm (top) and 20 µm (bottom).

**Figure 2.**
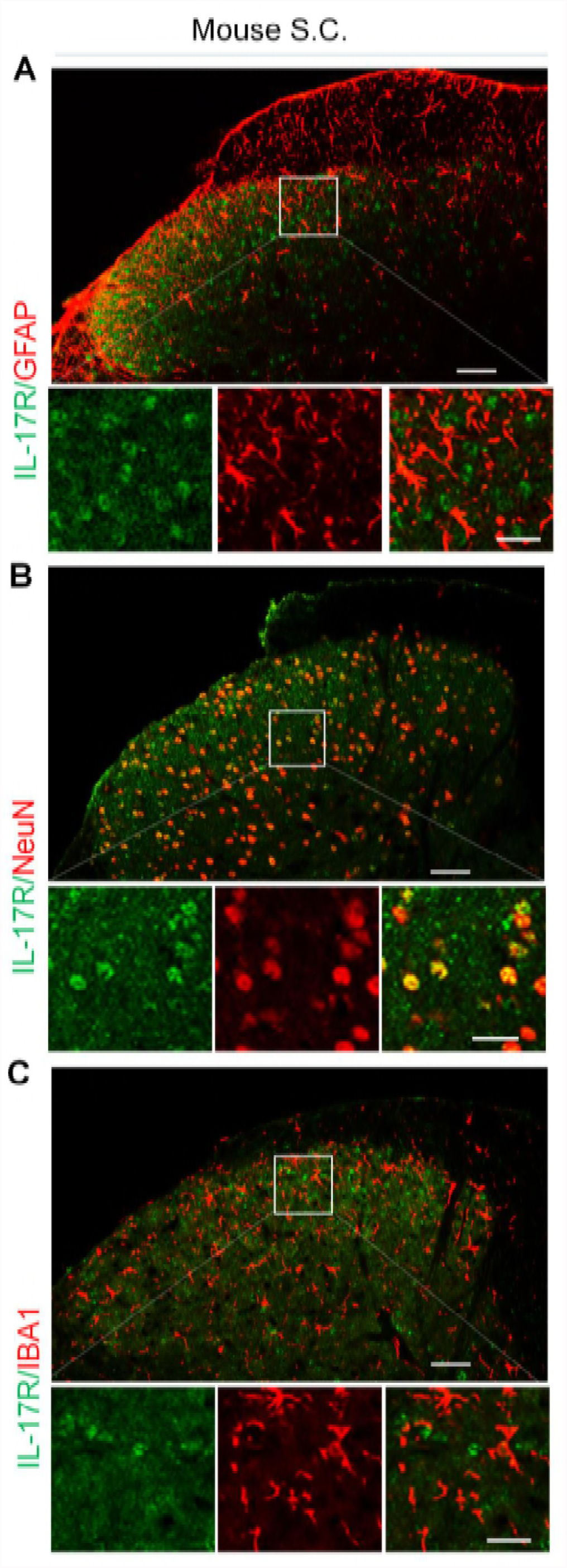
Photomicrographs showing IL-17 receptor (IL-17R) expression and colocalization of IL-17R with NeuN in SDH. Double labeling of IL-17R with astrocyte marker GFAP (**A**), neuron marker NeuN (**B**), and microglia marker IBA1 (**C**) in SDH. The white square in the top image is enlarged in three separate boxes with single and merged images in each picture (**A**, **B** and **C**). The scale bars represent 50 µm (top) and 20 µm (bottom).

### IL-17 enhances excitatory synaptic transmission and potentiates NMDA-mediated eEPSC in spinal cord slices

SOM^+^ neurons are excitatory interneurons and indispensable for mechanical pain (Duan et al., 2014a, Duan et al., 2018). These neurons also exhibit marked synaptic plasticity in pathological pain conditions (Xie et al., 2018, Xu et al., 2013). We first recorded spontaneous EPSCs (sEPSCs) in outer lamina II (II_o_) SOM^+^ neurons in isolated spinal cord slices of SOM-tdTomato mice (Fig. 3A). Acute perfusion of IL-17, at a low concentration (10 ng/mL, 3 min), induced a rapid and significant increase in 8 out of 10 neurons in the frequency of sEPSCs (Fig. 3 B, C, E, p=0.0003), suggesting a possible presynaptic mechanism of IL-17 to enhance glutamate releases. Notably, IL-17 produced a 57% increase in sEPSC frequency. Because excitatory synaptic transmission is mainly mediated by AMPA and NMDA receptors (AMPAR and NMDAR) and NMDAR is critical for spinal cord synaptic plasticity and pathogenesis of pain (Woolf and Salter, 2000), we further examined the effects of IL-17 on NMDAR-EPSC evoked by dorsal root entry zoon (LT) stimulation. The amplitude of NMDAR-EPSC was also significantly increased by IL-17 (Fig. 3 G, H, 37%, p=0.0077), suggesting a positive regulation of excitatory synaptic transmission by IL-17.

**Figure 3.**
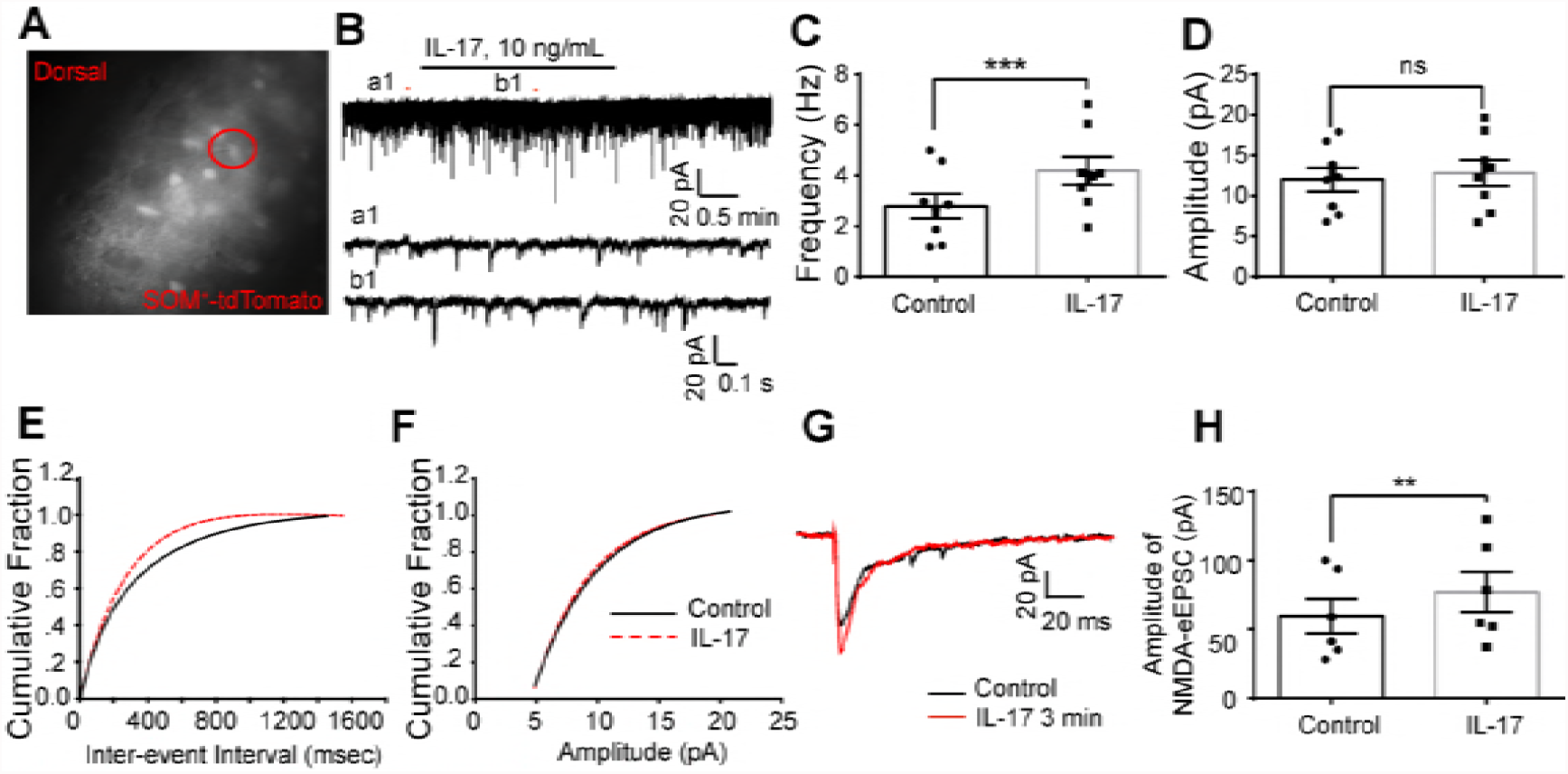
Potentiation of excitatory synaptic transmission by IL-17 in SDH lamina II_o_ SOM^+^ neurons. (**A**) Mouse spinal cord slice image showing a recording electrode in a SOM^+^ neuron (red circle). (**B**) Traces of sEPSCs in lamina II_o_ SOM^+^ neurons after perfusion of IL-17 (10 ng/mL, 2 min). a1 and b1 are enlargements of the recordings before and after IL-17 treatment, respectively. (**C**, **D**) Quantification of changes in frequency and amplitude of sEPSCs (n=8 neurons/group). (**E**, **F**) Corresponding cumulative distributions of inter-event interval and amplitude from one neuron. (**G**) Traces of NMDA-eEPSC before (black) and after (red) IL-17 treatment. (**H**) Potentiation of the amplitude of NMDA-eEPSC by IL-17 (n=6). ** P<0.01, ***P<0.001, two-tailed paired student’s test. ns, not significant. All the data were mean ± S.E.M.

### IL-17 decreases the inhibitory control of SOM^+^ neurons and suppresses GABA-induced currents

SOM^+^ excitatory neurons receive inhibitory input from inhibitory neuron (Duan et al., 2014b). We next recorded spontaneous IPSC (sIPSCs) in lamina II_o_ SOM^+^ neurons by using a pipette solution containing Cs^2+^. After exposure of spinal cord slice to IL-17 (10 ng/mL) for 3 min, most neurons (7 out 10) responded to IL-17. IL-17 produced a significant decrease of sIPSCs in both frequency (Fig. 4A, B, E, p=0.0039) and amplitude (Fig. 4C, F, p=0.0019). Because inhibitory synaptic transmission in the spinal cord is mediated by GABA and glycine, two major inhibitory neurotransmitters (Todd, 2010), we further assessed if IL-17 would also alter GABA and glycine evoked currents in lamina II_o_ SOM^+^ neurons. Bath application of GABA (100 µM) and glycine (1 mM) induce marked inward currents. Interestingly, acute application of IL-17 (10 ng/ml) only inhibited GABA-induced current (Fig. 4D, I, P=0.0016) but had not effect on glycine-induced current in spinal SOM^+^ neurons (Fig. 4H, J, P=0.2540), suggesting a specific regulation of IL-17 on GABAR-mediated inhibitory synaptic transmission.

**Figure 4.**
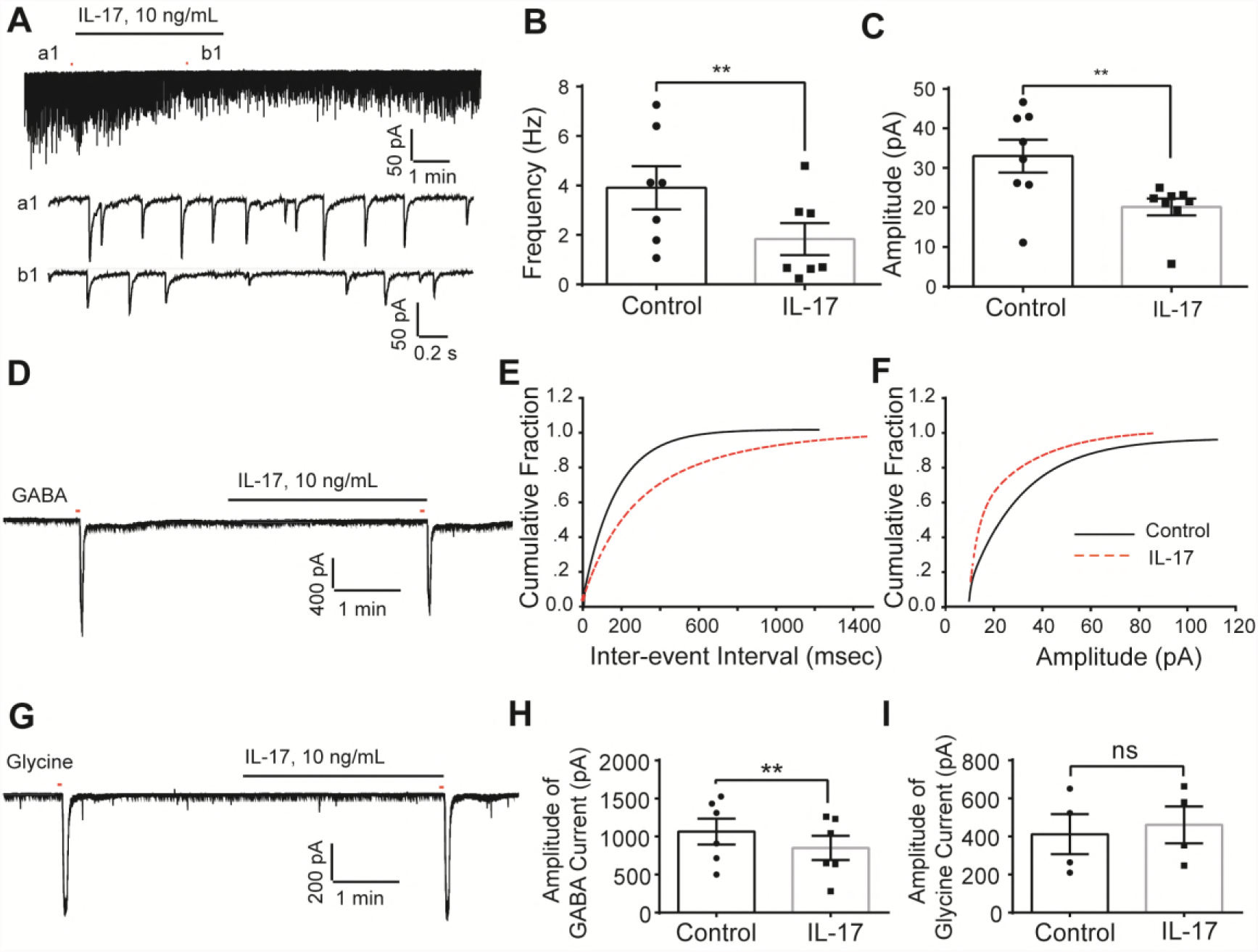
Suppression of inhibitory synaptic transmission in SOM^+^ excitatory neurons by IL-17 in spinal cord slices. (**A)** Typical traces of sIPSC in lamina II SOM^+^ neurons after perfusion of IL-17 (10 ng/mL, 2 min). a1 and b1 are enlargements of the recordings before and after IL-17 treatment, respectively. (**B**, **C**) Quantification of changes in frequency and amplitude of sIPSC (n=7, neurons/group). (**D**) Traces of GABA-induced current before (left) and after (right) IL-17 treatment. 100 μm GABA was applied for 3 seconds to induce an inward current. (**E**, **F**) Corresponding cumulative distributions of inter-event interval and amplitude from one neuron. (**G**) Traces of glycine-induced current before (left) and after (right) IL-17 treatment. 1 mM glycine was applied for 3 seconds to induce an inward current. (**H**) Suppression of the amplitude of GABA-induced current by IL-17 (n=6 neurons/group). (**I**) No changes in the amplitude of glycine-induced current by IL-17 (n=4).** P<0.01, two-tailed paired student’s test. ns, not significant. All the data were mean ± S.E.M.

### Up-regulation of endogenous IL-17 and IL-17R regulates synaptic plasticity in paclitaxel-treated mice

We first tested whether paclitaxel alters IL-17 levels in CSF and spinal cord. The CSF and spinal cord dorsal horns were collected from mice with confirmed mechanical allodynia at day 7 after paclitaxel treatment. ELISA analysis demonstrated significant increases in IL-17 levels in the spinal cord dorsal horn and CSF samples of paclitaxel-treated mice vs. vehicle-treated mice (Fig. 5A,B, p=0.0092). Also, we found that majority of spinal SOM^+^ neurons express IL-17R, providing an anatomical basis for IL-17 to directly regulate the activities of SOM^+^ neurons (Fig. 5 C).

**Figure 5.**
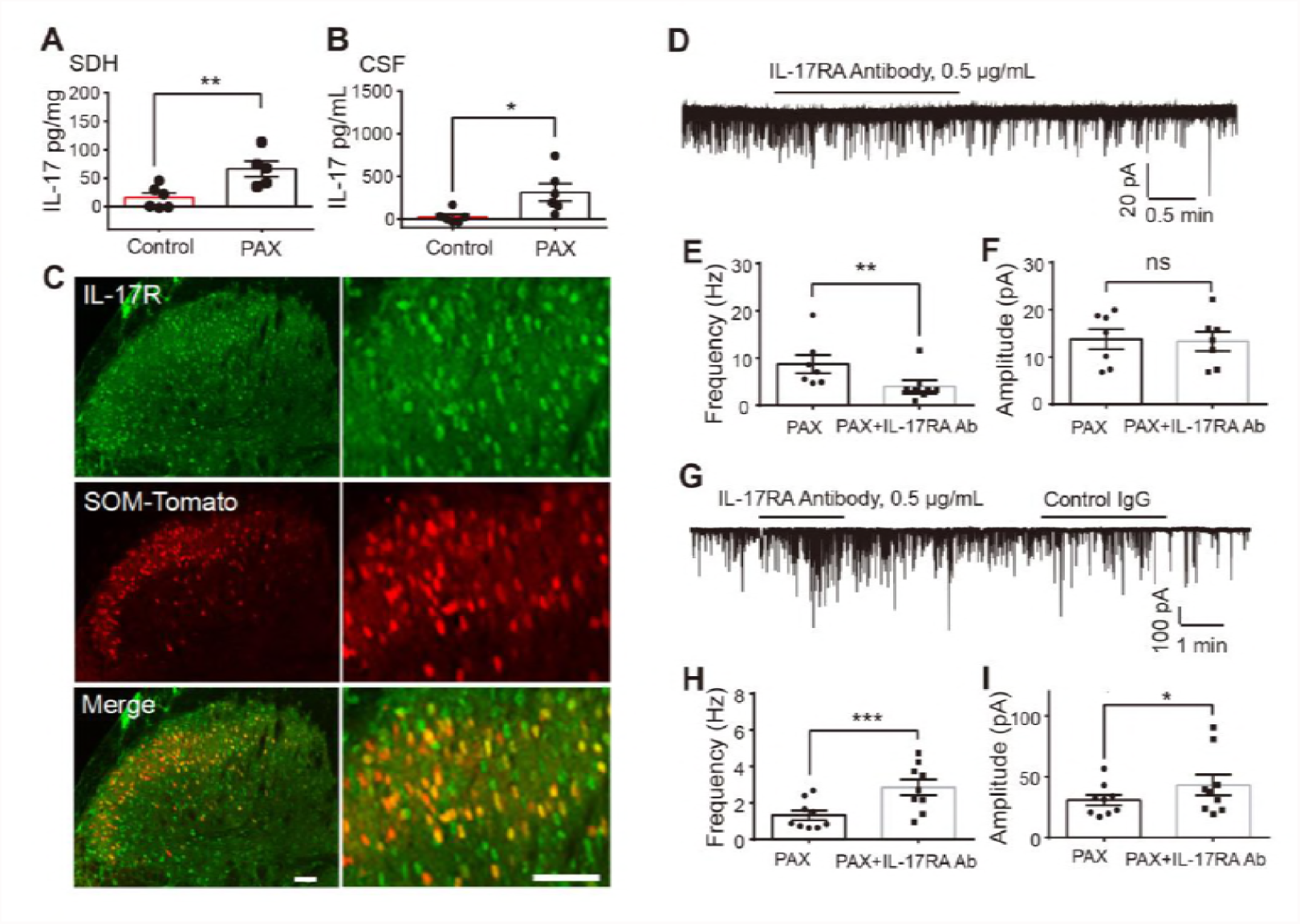
Endogenous IL-17 regulates synaptic transmission in paclitaxel-treated mice via IL-17R. (**A**, **B**) ELISA analysis showing IL-17 levels in SDH (**A)** and CSF (**B**) samples of control and paclitaxel treated mice. The samples were collected 7 d after the paclitaxel treatment. * P<0.05, ** P<0.01, two-tailed unpaired student’s test. n=6 animals/group. Sample sizes are indicated in each graph. (**C**) IL-17R expression in SOM^+^ neurons in SDH. Scale bar: 50 μm. (**D**) Traces of sEPSCs in lamina IIo SOM^+^ neurons following paclitaxel treatment before and after perfusion of IL-17 receptor A antibody (IL-17RA Ab, 0.5 μg/mL, 2 min). (**E**, **F**) Quantification of change in frequency and amplitude of sEPSCs (n=7). (**G**) Typical Traces of sIPSC in lamina II SOM^+^ neurons 7 days following paclitaxel treatment before and after perfusion of IL-17RA Ab (0.5 μg/mL, 2 min). (**H**, **I**) Quantification of changes in frequency and amplitude of sIPSC (n=9). * P<0.05, ** P<0.01, *** P<0.001, two-tailed paired student’s test. ns, not significant. All the data were mean ± S.E.M.

On the basis of these results, we postulated that IL-17 upregulation after chemotherapy contributes to spinal cord synaptic plasticity (i.e. central sensitization), a driving force of pathological pain (Ji, 2017). We measured the frequency and amplitude of sEPSCs or sIPSCs of spinal SOM^+^ neurons in paclitaxel-treated mice and tested the involvement of endogenous IL-17 in synaptic plasticity after CIPN. Blocking IL-17R with a neutralizing antibody resulted in opposite changes in excitatory and inhibitory synaptic transmission in lamina II_o_ SOM^+^ neurons of paclitaxel-treated animals: a decrease in frequency of sEPSCs (Fig. 5D, E, F, p=0.0019) but an increase in sIPSC frequency (Fig. 5G, H, p=0.0004) and sIPSC amplitude (Fig. 5I, p=0.0384) However, control IgG had no effects on sIPSC (Fig. 5G). Thus, endogenous IL-17 is involved in modulating excitatory and inhibitory synaptic transmission in spinal SOM^+^ neurons via IL-17R after paclitaxel treatment, suggesting a role of the IL-17-IL-17R pathway in CIPN.

### IL-17 increases the excitability of small-sized mouse and human DRG neurons

Immunostaining revealed that IL-17R is expressed by small-sized mouse DRG neurons, and some of them bind IB4 (Fig. 6A). This is consistent with a previous report that IL-17RA is localized in majority of rat DRG neurons (Richter et al., 2012). In contrast, IL-17 expression was observed in satellite glial cells that express glutamine synthetase (GS, Supplementary Figure 1). This respective expression of IL-17 and IL-17R in glia and neurons in the DRG is similar to that observed in the spinal cord, suggesting that IL-17 and IL-17R can mediate neuron-glial interactions both in the central and peripheral nervous system.

**Figure 6.**
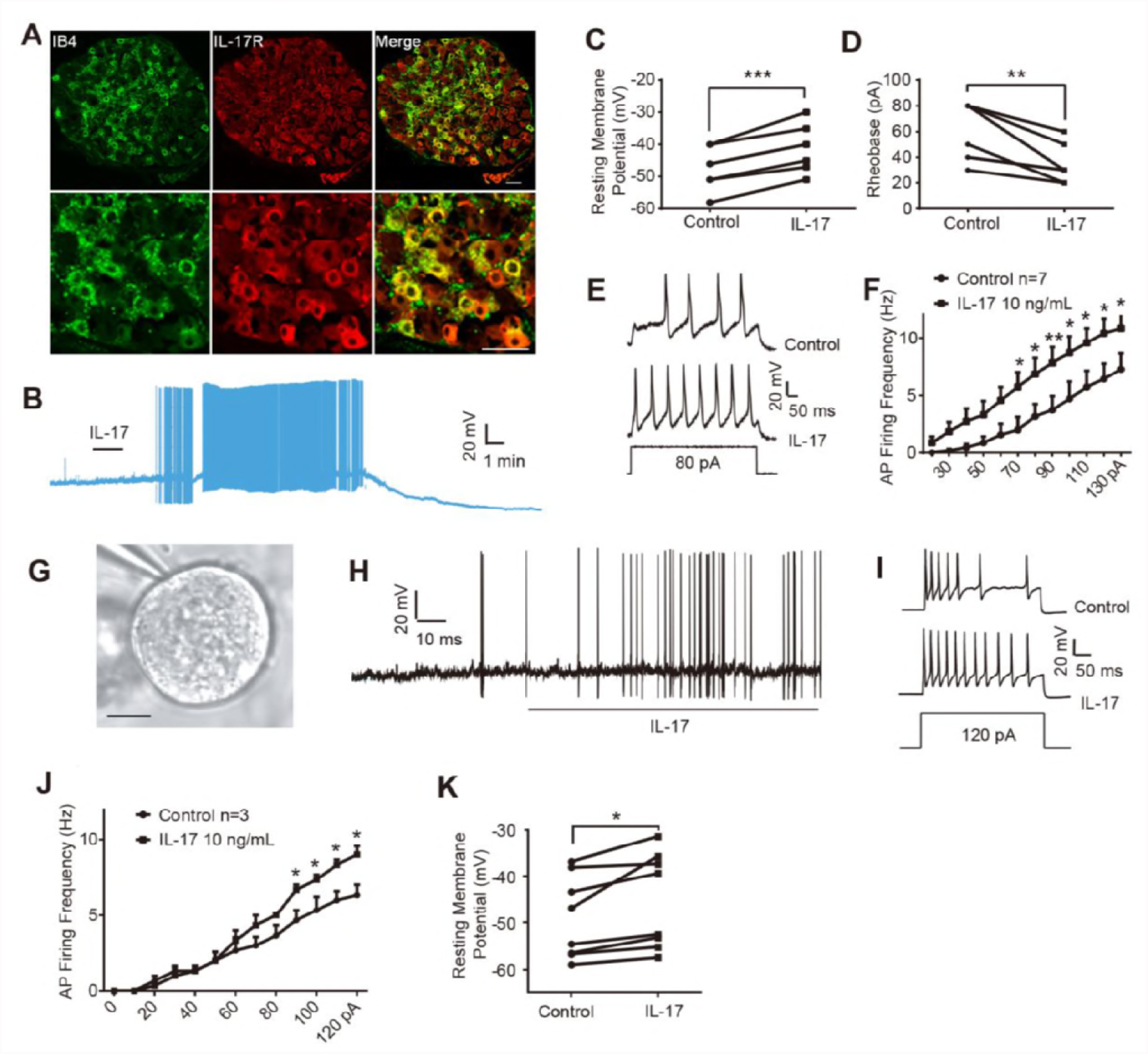
IL-17 increases the excitability of dissociated mouse DRG (mDRG) and human DRG neurons. (**A**) IL-17R expression in mDRG neurons. Scale bar: 50 μm. (**B**) Acute application of IL-17 (10 ng/mL) evoked spontaneous action potentials (APs) in mDRG neurons. (**C**, **D**) The effects of IL-17 on resting membrane potential (**C**) and rheobase (**D**). ** P<0.01, *** P<0.001, two-tailed paired student’s test; n = 6 neurons/group. (**E**) Traces of APs in small-sized mDRG neurons before and after perfusion of IL-17. (**F**) Quantification of firing frequency of action potentials as shown in **e**. * p < 0.05, ** P<0.01; two-way ANOVA; n = 7 neurons/group. (**G**-**J**) Whole-cell recording in dissociated small-diameter (<55 µm) human DRG neurons. (**G**) Image of an isolated human DRG neuron with the tip of a pipette during patch clamp recording. Scale bar: 20 μm. (**H**, **I**) The representative traces of action potentials. (**H**) Acute application of IL-17 (10 ng/mL) evoked spontaneous APs in a human DRG neurons. (**I**) Representative AP waveforms for the neuron (G) evoked by direct current injection before and after 2 min of acute perfusion with IL-17. (**J**) Quantification of firing frequency of action potentials. * P<0.05, ** P<0.01; two-way ANOVA; n = 3 neurons. (**K**) Quantification of RMPs before and after IL-17 treatment (10 ng/mL). *P <0.05, **P<0.01; paired t-test; n = 8 neurons.

To determine a role of IL-17 in regulating the excitability of DRG neurons, we tested the effects of IL-17 on dissociated small-sized mouse DRG neurons (< 25 µm in diameter) using whole-cell patch clamp recordings. Acute application of IL-17 (10 ng/ml) to mouse DRG neurons in vitro induced spontaneous discharge and bursts of action potentials in some DRG neurons (Fig. 6 B). Also, IL-17 significantly depolarized the resting membrane potential (Fig. 6 C, p=0.0001) and significantly decreased rheobase (Fig. 6 D, p=0.0068). IL-17 bath application also increased numbers of action potential discharges to suprathreshold current injection (Fig. 6 E, F). Therefore, IL-17 increases excitability of nociceptive neurons by altering rheobase and resting membrane potential in nociceptive neurons, leading to enhanced discharges of action potentials.

To enhance translational potential of this study, we also examined the action of IL-17 in human DRG neurons. An example of small-sized human DRG neuron from disease-free donors is shown in Figure 6G. Like mouse DRG neurons, human DRG neurons showed spontaneous action potentials and increased AP firing number during acute application of human IL-17 (10 ng/ml, Fig. 6 H-J). Also, IL-17 caused significant depolarization in the resting membrane potential on human DRG neurons (p = 0.016, Fig. 6K). These findings in isolated mouse and human DRG neurons show that IL-17 has the potential to directly increase the excitability of primary afferent neurons.

### IL-17R is required for hyperexcitability of mouse DRG neurons after paclitaxel chemotherapy

Paclitaxel was shown to increase responsiveness and excitability of mouse and human DRG neurons (Chang et al., 2017, Li et al., 2015b). We compared the number of APs evoked by a 600-ms current injection through intracellular electrode in mouse DRG neurons and tested the effects of paclitaxel after bath application (1 µM, 2 h). Only neurons that showed more than one AP to the stimulation were included in the analysis. Paclitaxel increased the AP firing numbers in small sized neurons compared with the vehicle-treated neurons (Fig. 7A, B). Strikingly, this excitability increase was suppressed by IL-17RA antibody, as compared with control IgG (Fig. 7 C, D). This result implies a direct regulation of neuronal hyperexcitability by IL-17R after chemotherapy.

**Figure 7.**
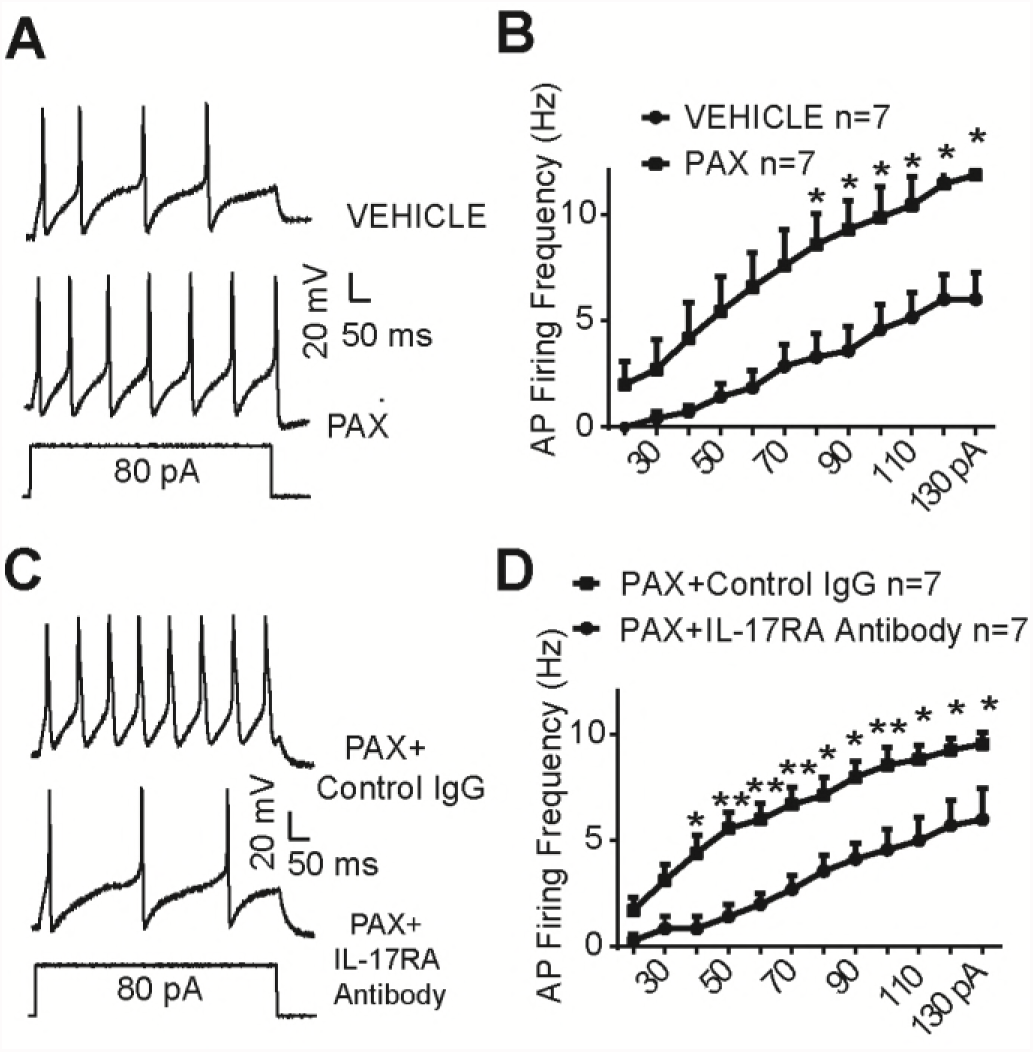
IL-17 Receptor A neutralizing antibody inhibits the hyperexcitability of small-sized mouse DRG (mDRG) neurons following paclitaxel treatment. (**A**) Traces of APs in mDRG neurons evoked by direct current injection pretreated with vehicle or 1 μM paclitaxel for 3 h. (**B**) Quantification of APs firing frequency pretreated with vehicle or paclitaxel. * p < 0.05; two-way ANOVA; and n = 7 neurons/group. (**C**) Traces of APs after application of Control IgG or IL-17RA Ab (0.5 μg/mL, 2 min) in paclitaxel pretreated mDRG neurons. (**D**) Quantification of APs firing frequency as shown in **D**. * p < 0.05, ** P<0.01; two-way ANOVA; and n = 7 neurons.

### IL-17 and IL-17R contribute to mechanical hypersensitivity after chemotherapy

To test a central role of IL-17 in pain modulation, we compared mechanical pain thresholds of mice following intrathecal injection of IL-17 and vehicle. Spinal injection of low dose of IL-17 (50 ng, i.t.) resulted in a transient reduction in paw withdrawal threshold (PWT) in 1 h, but the mechanical allodynia recovered in 2 h (Fig. 8A). A high dose of IL-17 (100 ng, i.t.) caused a more persistent reduction in PWT for 3 h, recovering after 5 h (Fig. 8A). These data suggest that IL-17 is sufficient to induce pain hypersensitivity in naïve animals.

**Figure 8.**
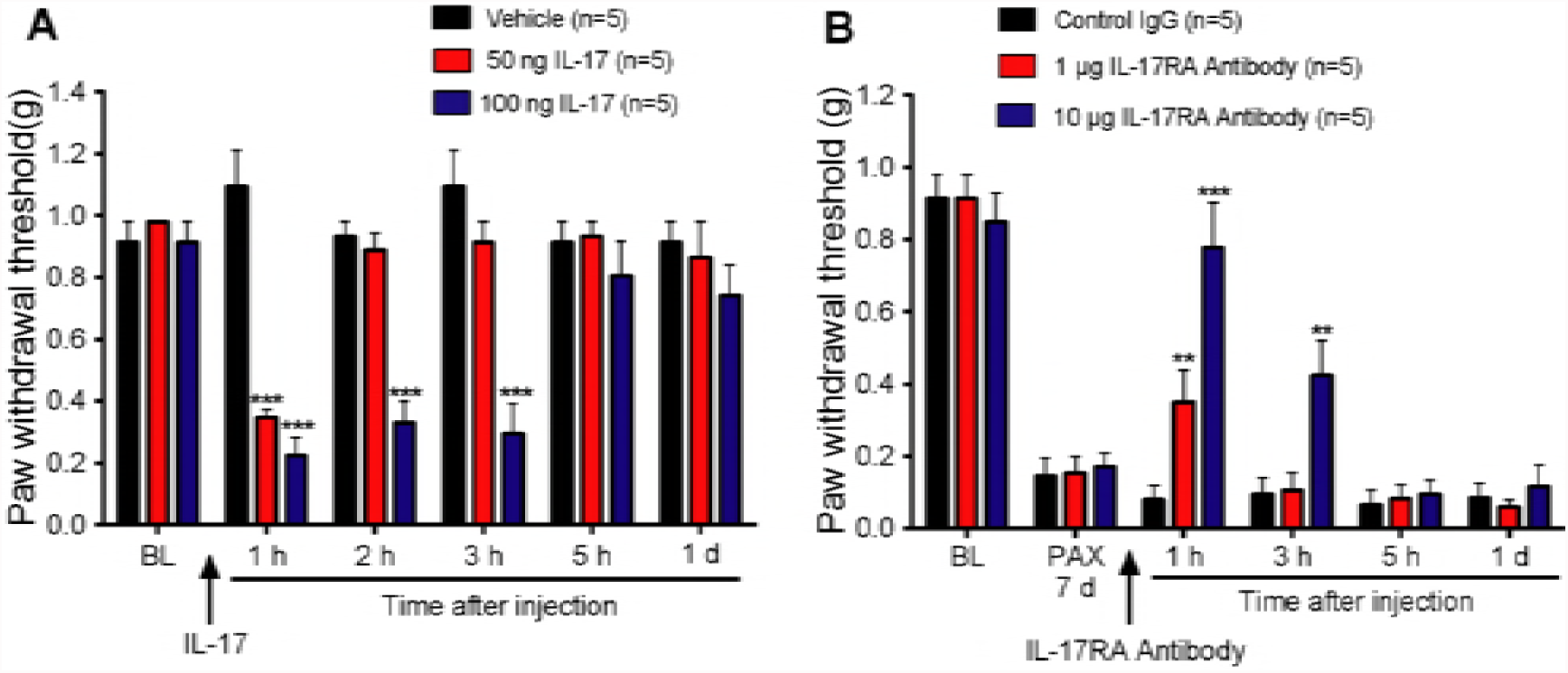
IL-17 and IL-17 receptor A (IL-17RA) contribute to paclitaxel-induced neuropathic pain. (**A**) Intrathecal injection of IL-17 induces a transient and dose-dependent mechanical allodynia (i.e. reduction in PWT). (**B**) Mechanical allodynia, induced by paclitaxel (6 mg/kg, i.p.), is attenuated by intrathecal injection of IL-17RA Antibody (1 and 10 µg). * P < 0.05; **P < 0.01; ***P< 0.001; vs. control IgG (10 µg, i.t.), two-way ANOVA; n = 5 mice/group.

Finally, we investigated the contribution of IL-17R to chemotherapy-evoked neuropathic pain. A single injection of paclitaxel (6 mg/kg, i.p.) evoked a remarkable reduction in PWT on day 7, which was reversed by IL-17RA antibody (10 µg, i.t.), in a dose-dependent manner. Intrathecal injection of control IgG (10 µg) produced no changes in PWT (Fig. 8B). The results indicate that IL-17R is required for maintaining chemotherapy-induced neuropathic pain.

## Discussion

We have provided new insights into how IL-17 promotes chemotherapy-induced neuropathic pain. Our results show that IL-17 and IL-17R regulate neuropathic pain via multiple mechanisms, including neuron-glial interactions, central sensitization, and peripheral sensitization (Supplemental Figures 1 and 2). In the spinal cord, IL-17 increases NMDA receptor-mediated currents and facilitates excitatory synaptic transmission. In particular, IL-17 suppresses inhibitory synaptic transmission by inhibiting GABA receptor-mediated currents. In the DRG, IL-17 increases neuronal excitability and IL-17R contributes to paclitaxel-induced nociceptor hyperactivity.

### IL-17 and IL-17R mediate neuron-glial interactions both in the central and peripheral nervous system

Recent progress has demonstrated critical roles of spinal glial cells in driving chronic pain via production of proinflammatory cytokines and neuron-glial interactions (Ji et al., 2016, McMahon and Malcangio, 2009, Ren and Dubner, 2010, Gosselin et al., 2010).

Microglia and astrocytes play different roles in different pain conditions, such as CIPN. Paclitaxel was shown to induce astrocyte activation but not microglia activation in the spinal cord (Zhang et al., 2012b). Activation of p38 MAP kinase in spinal microglia contributes to neuropathic pain after nerve trauma and cancer pain (Jin et al., 2003, Yang et al., 2015). However, spinal inhibition of p38 MAP kinase fails to affect chemotherapy-induced mechanical allodynia (Luo et al., 2017). Our results also highlight a role of astrocytes in CIPN. IL-17 is a T cell-derived cytokine, but we found IL-17 immunoreactivity exclusively in GFAP-expressing astrocytes. In contrast, IL-17R was primarily expressed in spinal cord neurons including SOM^+^ neurons. This unique localization of IL-17 and IL-17R offers a cellular basis for astroglia-neuro interaction in pain regulation. In parallel, we observed respective expression of IL-17 and IL-17R in satellite glial cells and neurons in mouse DRG. Thus, IL-17/IL-17R signaling could promote both central sensitization and peripheral sensitization via neuron-glial interactions in the CNS and PNS.

### IL-17 and IL-17R modulate excitatory synaptic transmission in the spinal cord pain circuit

Enhanced excitatory synaptic transmission has been shown in spinal cord neurons including SOM^+^ neurons in various pathological pain conditions (Chen et al., 2015, Yang et al., 2015). Our data indicates that IL-17 is both sufficient and required for inducing this synaptic plasticity. Exogenous IL-17 rapidly increased EPSC in spinal cord slices from naïve animals. Spinal cord slice from paclitaxel-treated animals exhibited an increase in EPSCs, which was suppressed by IL-17R antibody, suggesting an endogenous role of IL-17 in CIPN. Mechanistically, IL-17 acutely enhanced amplitude of NMDA evoked currents following dorsal root stimulation, suggesting that IL-17 increases NMDAR activity via rapid post-translational regulation. This is consistent with the previous report that IL-17 acts on spinal nociceptive neurons co-expressing IL-17R and NR1 to modulate pain (Meng et al., 2013). NMDAR plays a critical role in the induction and maintenance of central sensitization during persistent pain conditions (Woolf and Thompson, 1991, South et al., 2003). It remains to be investigated how IL-17 modulates NMDAR activity. It is possible that IL-17 activates protein kinases such as extracellular-regulated kinase (ERK) and protein kinase C to enhance NMDAR activation and neuronal excitability (Hu and Gereau, 2003). For example, TNF-α increases NMDA currents in spinal cord lamina II_o_ neurons via ERK phosphorylation (Xu et al., 2010). Thus, IL-17 plays a role in spinal pain modulation in part via NMDAR-mediated glutamatergic synaptic transmission.

### Modulation of inhibitory synaptic transmission by IL-17 and IL-17R

One of the most interesting findings of this study is profound suppression of inhibitory synaptic transmission in lamina II_o_ SOM^+^ neurons. Disinhibition, i.e., loss of inhibitory synaptic transmission is emerging as a key mechanism of neuropathic pain (Coull et al., 2003, Zeilhofer et al., 2012a, Lu et al., 2013). Removal of spinal inhibition, especially presynaptic GABAergic inhibition, not only reduces the fidelity of normal sensory processing but also provokes symptoms very much reminiscent of inflammatory and neuropathic chronic pain syndromes (Zeilhofer et al., 2012b, Takazawa et al., 2017, Chen et al., 2014). Our study shows that exogenous IL-17 rapidly (within a minute) and drastically decreased the frequency and amplitude of sIPSC. Mechanistically, IL-17 specially suppressed GABA but not glycine induced currents. Although TNF-α and IL-1β were also shown to regulate inhibitory synaptic transmission in spinal cord neurons (Kawasaki et al., 2008, Zhang et al., 2010, Chirila et al., 2014), they act on different pain circuits in the spinal cord. Multiple mechanisms have been implicated in disinhibition in pathological pain (Zeilhofer et al., 2012b). Our data suggest that IL-17 can elicit a very rapid loss of inhibition to open the spinal gate, which allows low-threshold mechanical stimuli to activate pain transmission neurons as predicted by the “Gate control theory” (Melzack and Wall, 1965, Wall, 1978). Majority of SOM^+^ excitatory neurons distribute in lamina II. These neurons not only receive an excitatory input from C-, A**β**- and A**δ**-fibers, but also receive an inhibitory control from inhibitory neurons (Duan et al., 2014b). Thus IL-17 may induce pain via distinct synaptic mechanisms either by increasing excitatory synaptic transmission or by decreasing inhibition control of SOM^+^ excitatory neurons. Our working hypothesis is illustrated in Supplementary Fig. 2

### Modulation of DRG neuronal excitability after chemotherapy

Paclitaxel (Taxol) is a widely used chemotherapeutic agent producing a neuropathy characterized by pronounced impairment of function in A-beta myelinated fibers, intermediate impairment of A-delta myelinated fibers, a relative sparing of C-fibers (Dougherty et al., 2004) and mechanical hypersensitivity (hyperalgesia and allodynia) (Polomano et al., 2001, Fossiez et al., 1996). Mechanical allodynia after paclitaxel is in part mediated by A-beta fibers (Xu et al., 2015). Paclitaxel increases excitability of DRG neurons via regulating the expression and function of ion channels such as TRPV1, TRPV4, HCN1, Nav1.7, leading to increased excitatory synaptic input to spinal cord SG neurons (Zhang and Dougherty, 2014, Li et al., 2015a, Chang et al., 2018). Peripheral mechanisms of pain modulation by IL-17 have been investigated. For example, IL-17 sensitizes joint nociceptors to mechanical stimuli to facilitate arthritic pain (Richter et al., 2012). It was also reported that neuronal IL-17R regulates mechanical but not thermal hyperalgesia by upregulation of TRPV4 but not TRPV1 in DRG neurons (Segond von Banchet et al., 2013). We observed rapid excitability increase in both mouse and human DRG neurons following IL-17 treatment, suggesting a possible post-translational modulation of some key ion channels, such as sodium channels. Our work in progress shows that IL-17 also increased sodium currents (data not shown). Interestingly, we found that the enhanced excitability in paclitaxel-pretreated small mouse DRG neurons can be abolished by a neutralizing antibody against IL-17R antibody. Since the recordings were conducted in dissociated neurons and physiological concentration of IL-17 is not present in culture medium, our result suggests a possibility that IL-17R may direct regulate neuronal activity in the absence of IL-17. Future study will examine how IL-17R interacts with ion channels such as Nav1.7. We should not rule out the possibility that satellite glial cells may also be attached to neurons in our culture conditions to communicate with neurons by releasing IL-17. Our working hypothesis of peripheral glial regulation of chemotherapy-evoked neuropathic pain via IL-17/IL-17R signaling is illustrated in Supplementary Fig. 3.

### Translational potential

IL-17 levels in sciatic nerves are elevated after nerve injuries (Noma et al., 2011). Our data show that IL-17 levels are also elevated in CSF and spinal cord in paclitaxel-treated mice. Importantly, intrathecal injection of IL-17 RA antibody effectively alleviated paclitaxel-induced neuropathic pain. Chemotherapy has been shown to activate cancer-associated fibroblasts to renewal cancer-initiating cells and maintain colorectal cancer by IL-17 secretion (Lotti et al., 2013). Thus, targeting IL-17 signaling may not only alleviate neuropathic pain but also improve anti-cancer efficacy after chemotherapy. The translational potential of this study is enhanced by demonstrating hyperexcitability of human sensory neurons in response to IL-17. IL-17 blockers have been developed for treating inflammatory diseases such as psoriasis and arthritis (Kivelevitch and Menter, 2015). Brodalumab (Kyntheum®) is a human anti-interleukin-17 receptor A (IL-17RA) monoclonal antibody available for use in patients with moderate to severe plaque psoriasis (Blair, 2018). Since mouse IL-17RA antibody is effective in suppressing neuronal hyperexcitability after paclitaxel, Brodalumab could be used to treat CIPN and neuropathic pain.

## Acknowledgments

This work was supported by grants from National Key R&D Program of China (2017YFB0403803), National Natural Science Foundation of China (31420103903, 31771164 and 81471130), Development Project of Shanghai Peak Disciplines Integrated Chinese and Western Medicine, and NIN R01 grants DE17794, DE22743, and NS87988 to R.R.J.

---

**Supplementary Figure 1.**
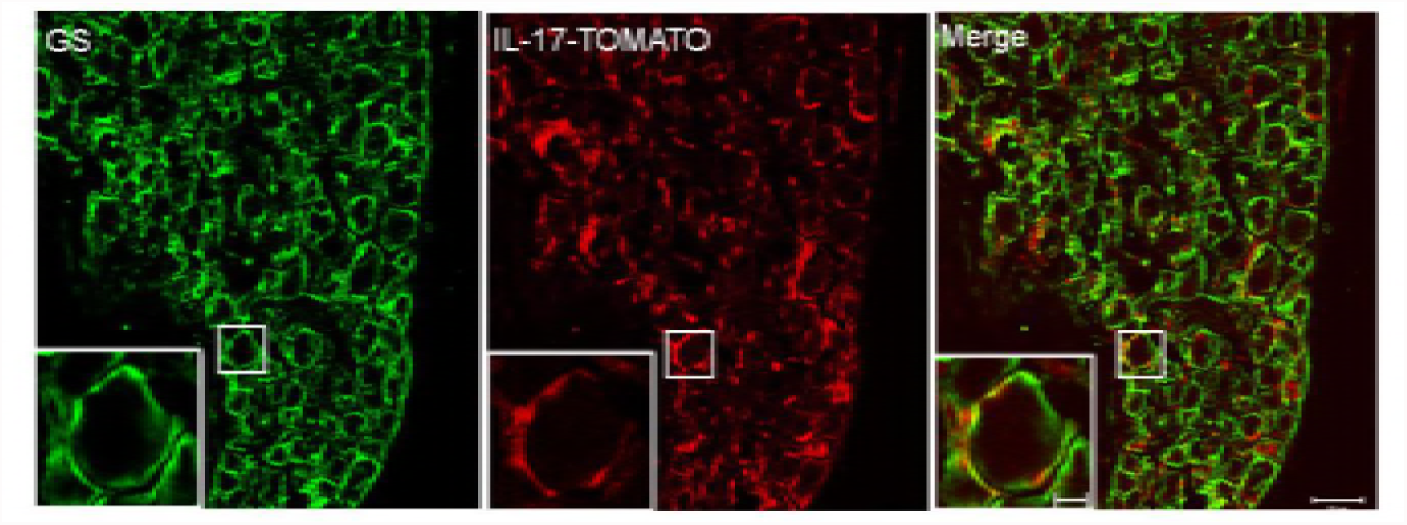
S1. Colocalization of IL-17 (red) with satellite glial marker glutamine synthetase (GS, green) Scale bar: 50 μm. The inserts are enlarged images. Scale bar: 10 μm.

**Supplementary Figure 2.**
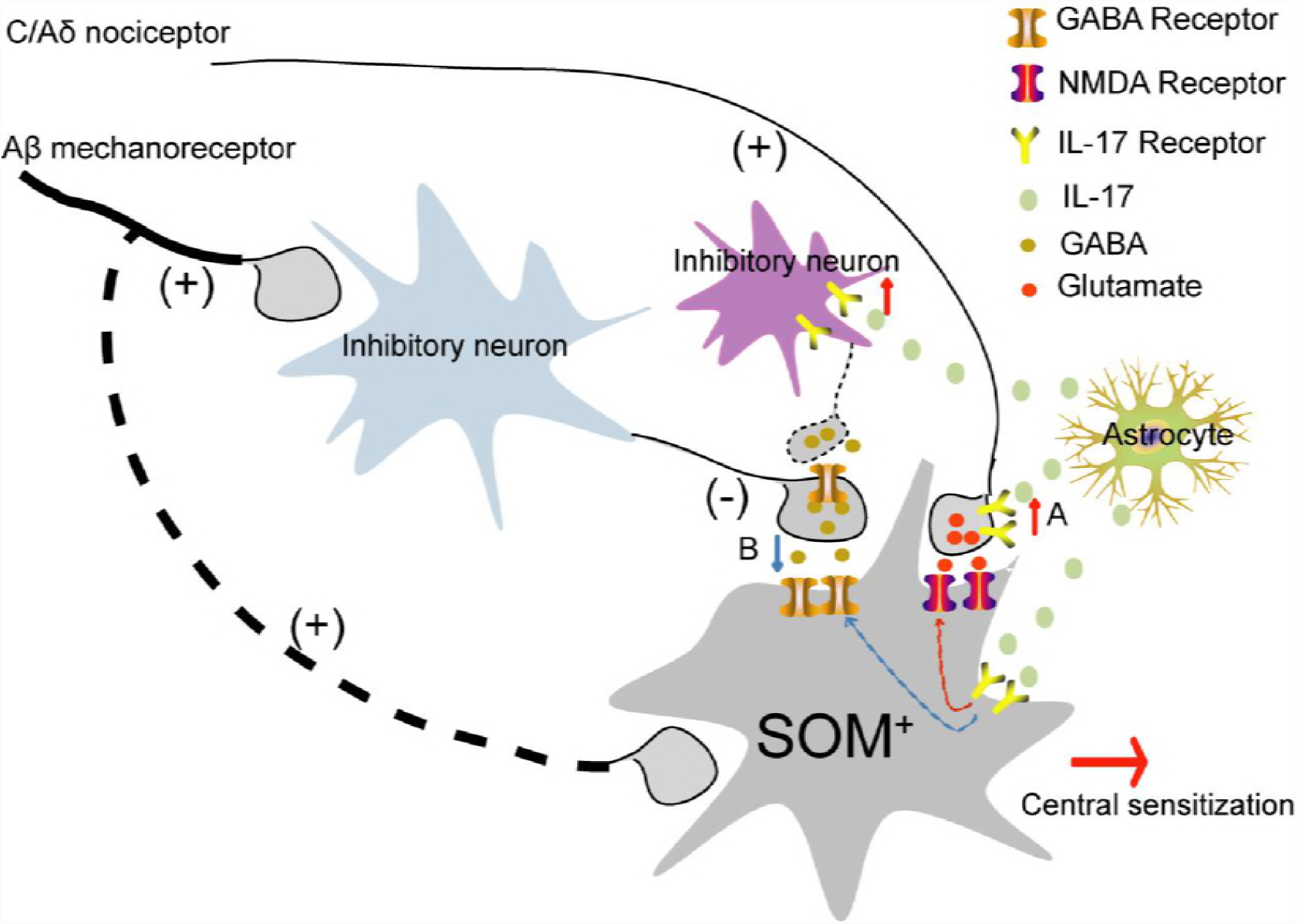
S2. A schematic showing how IL-17 modulates excitatory synaptic transmission (A) and inhibitory synaptic transmission (B) in spinal SOM^+^ neurons to induce central sensitization and pain hypersensitivity. (**A**) IL-17 is released from activated astrocytes and acts on IL-17R on presynaptic terminals and postsynaptic SOM^+^. Activation of presynaptic IL-17R results in increased glutamate release, whereas activation of postsynaptic IL-17R also enhances NMDAR activity. (**B**) IL-17 is also expressed by inhibitory neurons and facilitates GABA release to inhibit an inhibitory neuron, leading to a reduction in presynaptic inhibitory control of spinal SOM^+^ neurons. This disinhibition will also cause hyperactivity in SOM^+^ neurons.

**Supplementary Figure 3.**
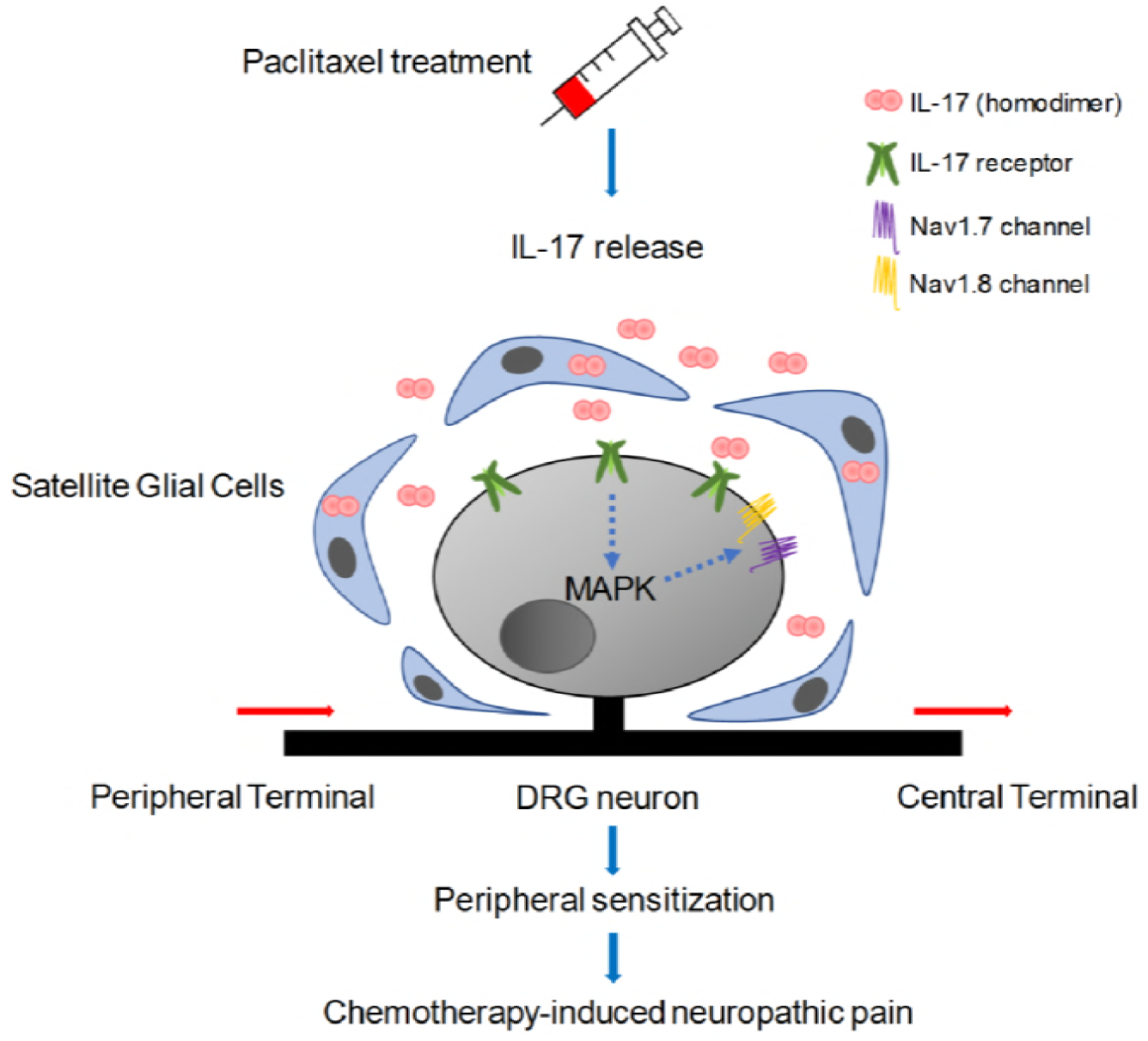
S3. A schematic showing IL-17-induced neuron-glial interaction in DRG. IL-17 is expressed by satellite glial cells. Following chemotherapy, IL-17 released from satellite glia acts on IL-17R on nociceptive neurons to increase neuronal excitability and induce peripheral sensitization.

